# Shifts from non-obligate generalists to obligate specialists in simulations of mutualistic network assembly

**DOI:** 10.1101/2022.08.04.502756

**Authors:** Timo Metz, Nico Blüthgen, Barbara Drossel

## Abstract

Understanding ecosystem recovery after perturbation is crucial for ecosystem conservation. Mutualisms contribute key functions for plants such as pollination and seed dispersal. We modelled the assembly of mutualistic networks based on trait matching between plants and their animal partners that have different degrees of specialization on plant traits. Additionally, we addressed the role of non-obligate animal mutualists, including facultative mutualists or non-resident species that have their main resources outside the target site. Our computer simulations show that non-obligate animals facilitate network assembly during the early stages, furthering colonization by an increase in niche space and reduced competition. While non-obligate and generalist animals provide most of the fitness benefits to plants in the early stages of the assembly, obligate and specialist animals dominate at the end of the assembly. Our results thus demonstrate the combined occurrence of shifts from diet, trait, and habitat generalists to more specialised animals.

## 1 Introduction

While ecosystems are degraded or destroyed at a high rate due to habitat conversion, they are also able to recover on abandoned land (Chazdon, 2014; FAO, 2020; Poorter *et al*., 2021). Such secondary succession of communities and ecosystems is globally widespread today and depends on the availability of species pools in the surrounding landscape, much like primary succession of new islands or habitats. Recovering ecosystems, such as secondary forests, play an increasingly important role for biodiversity conservation (Chazdon, 2014; Moreno-Mateos *et al*., 2020; Strassburg *et al*., 2020). Therefore, it is crucial to understand the process of ecosystem recovery after disturbance. Traditionally, the recovery of ecosystems has been measured through rather simple parameters, such as species richness. This, however, disregards much of an ecosystem’s complexity, which is not only based on the presence of species, but also on their interactions (Moreno-Mateos *et al*., 2020). Mutualistic interactions are ubiquitous in nature and are important for sustaining terrestrial biodiversity. In mutualism, species provide a service or a resource to other species with little costs to themselves (Bronstein, 2015). In terrestrial ecosystems, it is often an animal guild which provides a service (such as seed dispersal, pollination or protection) and a plant guild which provides resources (such as fruits, nectar or pollen) (Bronstein, 2021). Mutualisms can either be obligate, whenever a population goes extinct without its mutualistic partners, or facultative, when a population is also able to survive on its own (Bronstein, 2015). The dependence of many species on mutualistic interactions makes the investigation of these interactions especially important in the context of ecosystem recovery. Many plant species are obligate mutualists as they depend highly on animal services for reproduction and establishment (Chazdon, 2014). For pollination mutualism, it is estimated that 87.5 % of flowering plants globally and 94% of flowering plants in the tropics are animal pollinated (Ollerton *et al*., 2011). For seed-dispersal mutualism, animals are important dispersal agents for a considerable proportion of plants, in the case of tropical rainforests for around 90% (Jordano *et al*., 2014).

The ability of animal mutualists to successfully colonize a recovering site depends on the vegetation structure (e.g. due to the availability of nesting, food and shelter sites) which changes during succession. Animal mutualist pioneers therefore are often highly mobile, flying taxa (Chazdon, 2014; Öckinger *et al*., 2018; Woodcock *et al*., 2012), which can provide reproductive services (pollination or seed dispersal) to plants in the regenerating site but rely also on resources that can be found in the surrounding environment (Dunning *et al*., 1992). For instance, frugivorous bats may defaecate seeds during flight (Gorchov *et al*., 1993; Hodgkison *et al*., 2003; Muscarella & Fleming, 2007); birds that are diet and habitat generalists perch in recovering sites, depositing seeds and speeding up recovery Carlo & Morales (2016); Carlo & Yang (2011). Other authors also find that many animal pioneers are generalists with respect to habitat (Bowman *et al*., 1990; Chazdon, 2014; Chazdon *et al*., 2009; Liebsch *et al*., 2008; Pinotti *et al*., 2015) and diet (May, 1982). Furthermore, studies with pollinating bees suggested that animal pioneers are generalists with respect to the traits of their mutualistic partners (Chazdon, 2014). While animal mutualists fulfilling these different concepts of generalism dominate early stages of recovery, animal species with more specialized requirements for habitat, diet, or specific mutualistic partners are expected to increase in number with time (Bowman *et al*., 1990; Chazdon, 2014; May, 1982; Pinotti *et al*., 2015).

A way to understand the emergence of structural features in networks, such as the degree of specialization, is the theoretical study of network assembly, usually by computer simulations, which has in the past largely focused on food webs (Valdovinos, 2019). Such assembly simulations are interpreted as modelling the recovery of a destroyed habitat or the formation of a new habitat, for instance when a new volcanic island emerges from the ocean. Colonization of a destroyed or new habitat is modelled by immigration of species from a species pool, and the colonization of the habitat is in these models due to successful immigration events.

To our knowledge, Campbell *et al*. (2011) introduced the first model to assemble mutualistic plant-pollinator networks. This model is based on trait matching, with the survival of a species being determined by the number of beneficial versus the number of detrimental links it has. The focus of that study is on the diversity of assembly trajectories and outcomes. A recently introduced model by Becker *et al*. (2022) adds realism by including a niche width for animals and by determining species survival based on explicit simulation of population dynamics, including demographic noise. The assembly process in this model leads to networks where specialists and generalists coexist. However, since all species of the model are obligate mutualists, the initial stages of the assembly process are very slow.

In this paper we study a model that includes mutualistic animals that are not necessarily dependent on the resources provided by the mutualistic partner plants in the recovering site. These could represent highly mobile non-resident animals such as birds or bats that only occasionally forage in or fly over a recovering site as they rely also on adjacent undisturbed habitats for nesting or foraging. The model would also suit facultative mutualists that do not feed exclusively on the resources provided by their plant partner, or capital breeders that live in the larval stadium as herbivores and can feed in their adult stadium on nectar to increase reproduction (Davis *et al*., 2016). We use in the following the term “non-obligate” mutualists for these species. Other features of the model are similar to those of (Becker *et al*., 2022), in particular all plant species in our model are obligate mutualists.

With our model, we will identify the influence of non-obligate animal mutualists on network assembly, especially during the early stages when only few resources and mutualistic partners are available in the recovering site. We investigate the idea that animal pioneers with a high degree of generalism can play a key role during assembly. Furthermore, we explore how their influence changes during network assembly as species richness increases. At last, we study how the non-obligate animal mutualists affect the time evolution of the structural features of the network, in particular the degree of specialisation of the obligate animal mutualists. Taken together, our results will provide a unified view and show the combined action of the observed trends from more generalist animals (with respect to habitat, diet, or trait spectrum of their plant partners) to more specialist animals.

## 2 Materials and Methods

The building blocks of our model are trait-based network assembly and deterministic as well as stochastic population dynamics for plants and animals that interact in a service-for-resource mutualism.

### 2.1 Trait-based network architecture

Each plant and each animal in the network is assigned a trait 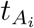 (for animals) and 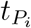 (for plants), and in case of animals also a niche width *s*_*j*_. We assume that this trait represents all the animal and plant properties that determine interaction and competition between species. The scale of trait values is chosen such that they lie in the interval [0,1]. Animal niche widths are chosen from the interval (0, 0.275], such that the model includes specialists with a narrow niche width as well as generalists that cover a considerable part of the trait space. Plants have a fixed niche width of *ξ* = 0.1 that determines their competition. The interaction between animals and plants is obtained from trait matching according to

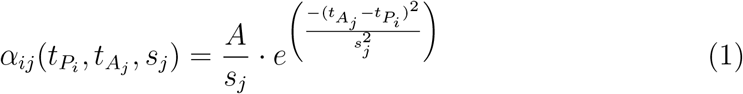

with 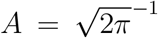, so that the area under the function is normalized to 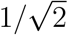. Species *P*_*i*_ and *A*_*j*_ interact only if *α*_*ij*_ *>* 0.05. According to Eq. (1), generalist animals with a high value of *s*_*j*_ (broad niche width) interact weakly, while specialist animals with a low value of *s*_*j*_ (narrow niche width) interact strongly.

### 2.2 Population dynamics equations

The dynamics for the population densities of plants *p*_*i*_ and animals *a*_*j*_ are given by the equations

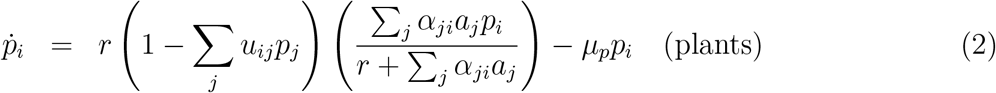

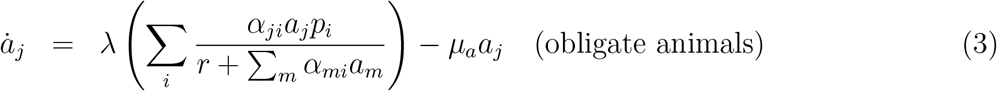

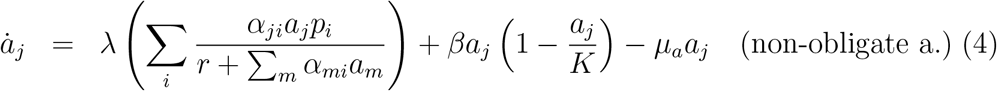

with the maximum plant growth rate *r*, an energy conversion efficiency of animals *λ/r*, the mortality rates of animals *µ*_*a*_ and plants *µ*_*p*_. For the non-obligate animals, there is a logistic growth term with the growth rate *β* and the carrying capacity *K*. This growth term makes sure that these animals are able to grow also in absence of a suitable mutualistic partner in the network (i.e., if all *p*_*i*_ = 0). The first two of these equations have been derived from a more extensive model that includes an equation for resource dynamics (Revilla, 2015), and they have also been used by Becker *et al*. (2022).

The plant competition coefficient *u*_*ij*_ is given by

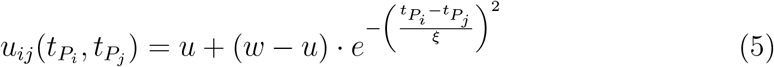

with the intraspecific competition strength *w*, the interspecific competition strength *u*, and the plant niche width *ξ*. For animals, competition for plant provided resources is implemented implicitly by a limitation of the total growth rate of all animals,

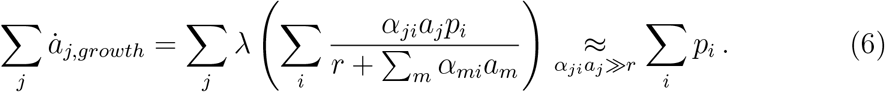

The cumulative animal population growth rate is of the order of the cumulative population density of plants, which in turn is proportional to the total amount of nectar or fruits.

Obligate animals and plants are assigned extinction thresholds *θ*_*a*_ and *θ*_*p*_, respectively. If the population density of a population falls below this threshold during the simulation, it is removed from the network.

These population dynamics equations can be used for any type of mutualism where plants provide resources and animals provide a service necessary for plant population growth.

### 2.3 Adding demographic noise to the population dynamics

Additionally to deterministic population dynamics, plant and pollinator population densities are subject to demographic noise, which is particularly relevant for small population sizes. The change in population sizes during the small time increment *dt* then becomes

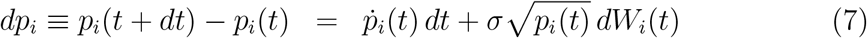

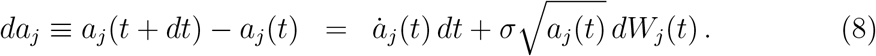

The strength of demographic noise is set via *σ*, which is 0 in the deterministic case and 0.1 in the simulations with demographic noise. *dW* is a random number that is taken from a normal distribution with mean 0 and standard deviation (*dt*)^1*/*2^. We scale the demographic noise with the square root of the population density because theoretical calculations of demographic noise through stochastic birth and death processes typically show that the variation coefficient of the population density scales with the inverse of the square root of the population density itself (Kot, 2001). In the computer simulations, time evolution was calculated using the Euler-Maruyama method (Higham, 2001), which is a method to numerically integrate stochastic equations. The time increment *dt* was set to 0.1 for all simulations, which is small compared to the time interval between immigration events.

### 2.4 Assembly simulations

The main aim of this study is to assess the influence of non-obligate animal populations on mutualistic network assembly. Therefore, we compare two versions of the same assembly model, namely one version with non-obligate animals, the other version without them. In the first version, we include a fixed number of non-obligate animal species from the beginning, because we assume that they are able to visit and forage in a recovering habitat even in the absence of mutualistic plant partners. During the course of the simulation, for both versions, obligate animals and plants were added to the network with equal probability at a rate *µ*. They were assigned a small initial population density just above the respective extinction thresholds *θ*_*p*_ and *θ*_*a*_, and a random trait and niche width, which are taken from a uniform distribution. The assembly was run until the long-term behavior of the system became visible.

For the model version with non-obligate animals, we initiated the assembly with 5 non-obligate animals with random traits and niche widths. This number is sufficient to cover a considerable part of the trait space, but still not too large to competitively exclude obligate animal species from entering. As starting point of the assembly, we added one plant species that was capable of surviving in the presence of the non-obligate animal species and had an initial population density of 5*·θ*_*p*_. The initial population density of the non-obligate animals was set to the carrying capacity *K*, which was 0.2 or 2 for different sets of runs. The two different values were chosen because a value of 0.2 leads to significantly lower and a value of 2 to similar population densities of non-obligate animals compared to obligate animals. No new non-obligate animals were introduced during the course of the simulation, and non-obligate animals could not go extinct since we assume that their populations are sustained by the surrounding unperturbed environment (or by other factors not included explicitly in the model).

For the simulations with only obligate mutualists, we generated initially one plant-animal pair with random traits and niche widths (provided they had a sufficiently strong interaction strength of *α*_*ij*_ *>* 3) to start off the network assembly. In this way, we accelerated the first step of the assembly, which consists in waiting for the almost simultaneous immigration of a pair that can interact with each other. The initial population densities were 5*·θ*_*p*_ (plants) and 10*·θ*_*a*_ (animals). In the rare case that all obligate animals and plants went extinct during the course of the assembly process, the assembly was restarted and the time counter was reset to 0.

### 2.5 Data analyses and evaluated quantities

For every parameter set, we calculated the number of plant and animal species in the network over time as well as the cumulative population densities of plants and animals (that is the sum over all population densities of plants and animals currently in the network at that specific time). We averaged over 100 simulation runs. Furthermore, we quantified the immigration success by keeping track of the proportion of immigrants with initial growth rate *>* 0 after immigration according to Eq. (2) and (3) and took averages over 500 simulation runs. To assess the importance of non-obligate animal mutualists, we evaluated the reproductive service provided by them. This was done by first evaluating their contribution to the growth term of each plant separately

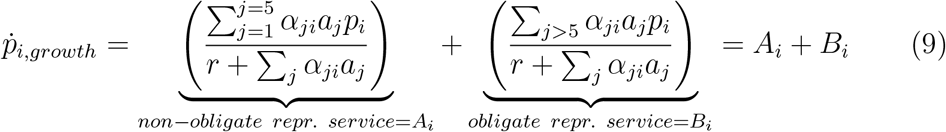

and then evaluating the relative contribution of non-obligate animals to the cumulative growth of all plants:

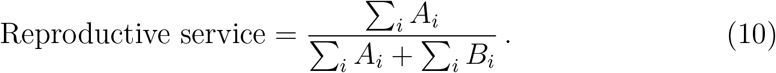

Additionally, we evaluated the reproductive service provided only by the most generalized and most specialized animals (lowest and highest 18% of the niche width interval) of each class (obligate or non-obligate).

The results displayed below are averages over 100 simulation runs.

## 3 Results

### 3.1 Communities assemble faster with non-obligate animal mutualists

Compared to the assembly simulations with only obligate mutualists, the simulations with non-obligate animals show a much faster increase of cumulative population densities and of species richness for both plants and obligate animals, see Fig. 1. The generality of this trend is confirmed by the fact that there is not much difference between the simulations with high and low carrying capacities (and thereby population densities) of non-obligate animals (compare the upper and middle rows of Fig. 1). The overall trends are similar for both model versions and agree with the findings reported in Becker *et al*. (2022): While the cumulative population densities of animals and plants increase monotonously from an initial value of 0 to a constant, maximum value for all investigated parameter sets, the species richness of obligate animal mutualists and of plants shows a peak at intermediate assembly stages and then decreases slightly towards its asymptotic value (see Fig. 1 upper and middle row) when population dynamics is deterministic. In the presence of demographic noise, the intermediate peak of species richness vanishes for the chosen parameter values (see Fig. 1 last row).

**Figure 1:**
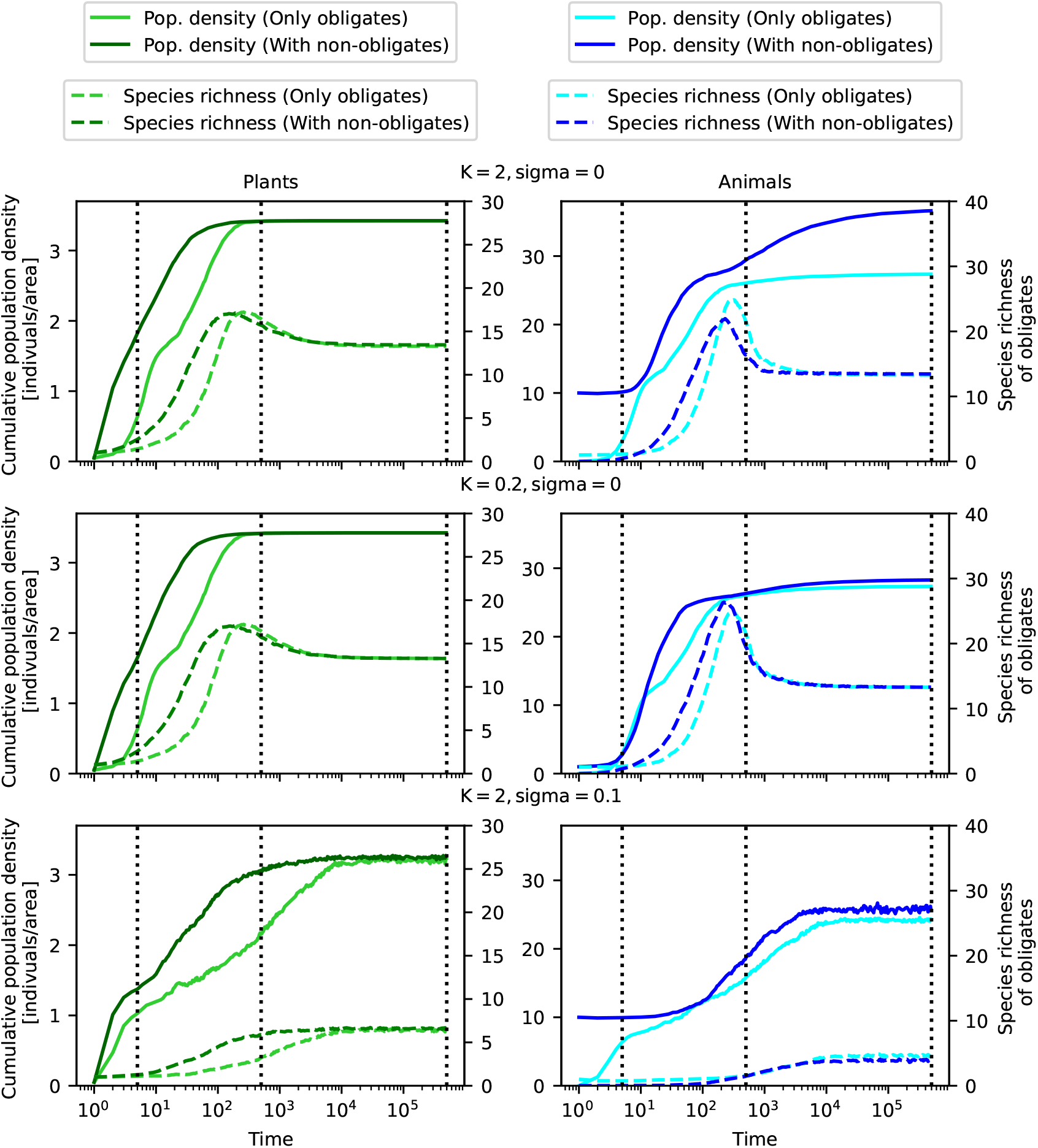
Cumulative population density and species richness of plants (left) and animals (right) over time for deterministic population dynamics (*σ* = 0) and population dynamics with demographic noise (*σ* = 0.1), and for high (*K* = 2) and low (*K* = 0.2) carrying capacity of non-obligate animals. Full lines correspond to cumulative population density (left y-axis), dashed lines to species richness (right y-axis). For species richness, only obligate animal and plant numbers are shown. For the cumulative population density, obligate and non-obligate animals are pooled together. Computer simulations of the model with only obligate animals (lighter colour) and the model with additional non-obligate animals (darker colour) are shown together. Vertical black dotted lines refer to times when network plots of Fig. 3 were created. The immigration rate was set to *µ* = 0.1.

Total animal cumulative population densities in the model with non-obligate animal mutualists are higher than in the model with only obligate mutualists. Species richness of obligate animals at the maximum is lower in the model with non-obligate animal mutualists compared to the model with only obligate animal mutualists. However, at the final stage of the assembly, we find similar numbers of obligate animal species in both model versions.

### 3.2 Immigration success of both mutualistic partners is enhanced by the presence of non-obligate animal mutualists

The faster increase of obligate species richness and total population densities in the simulations with non-obligate animals is caused by a higher immigration success in the presence of non-obligate animals, see Fig. 2. This increase in immigration success is higher for plants than for animals. Again, there is little difference between the simulations with high and low carrying capacities (i.e. population densities) of non-obligate animals. At later assembly stages, the immigration success decreases towards zero in the simulations with deterministic population dynamics, but it stabilizes at a non-zero value in the presence of demographic noise (see Fig. 2. This asymptotic value is affected only very weakly by the presence of non-obligate animals.

**Figure 2:**
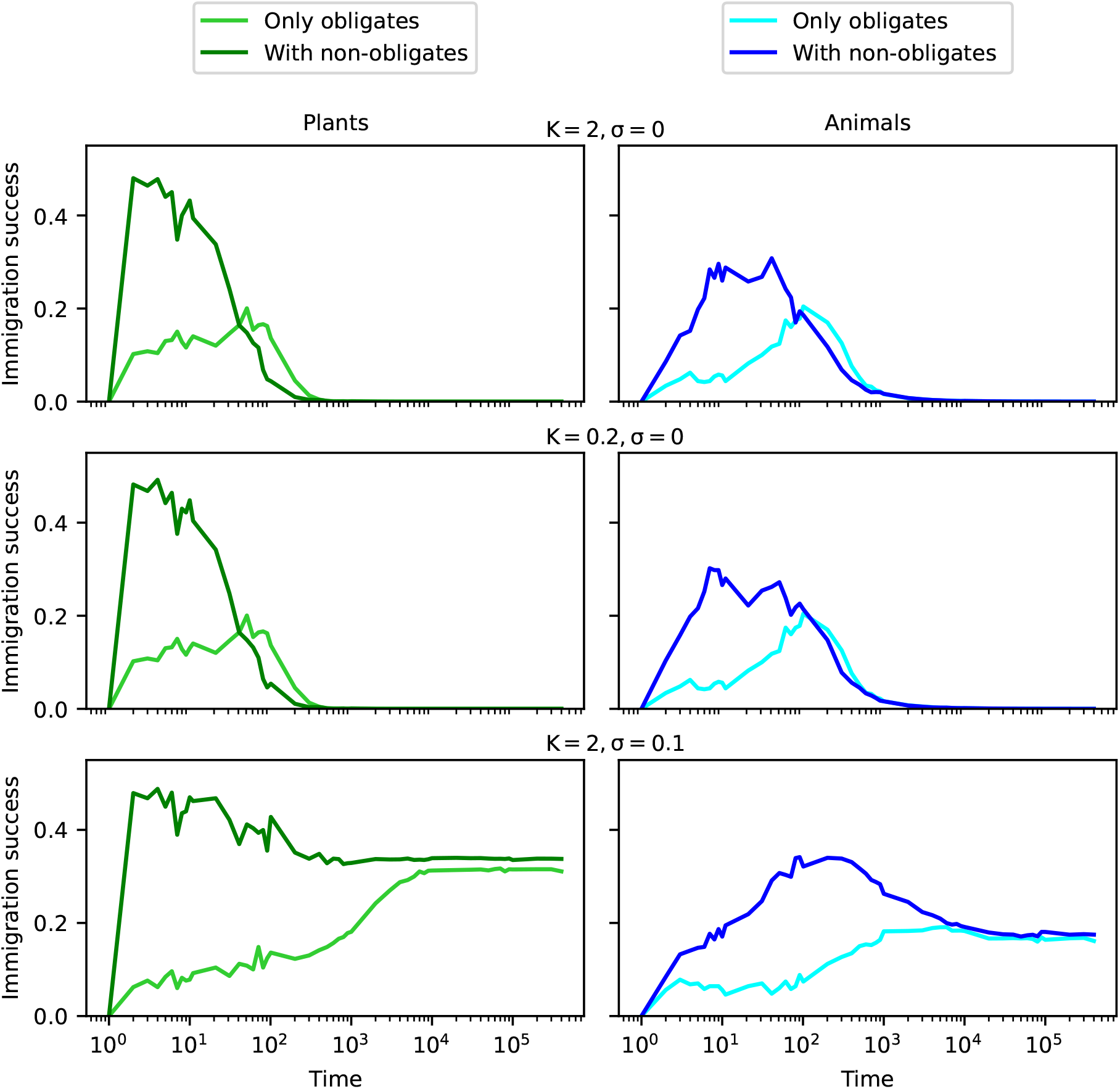
Immigration success of plants (left) and animals (right) over time for deterministic population dynamics (*σ* = 0) and population dynamics with demographic noise (*σ* = 0.1), and for high (*K* = 2) and low (*K* = 0.2) carrying capacity of non-obligate animals. Computer simulations of the model with only obligate animals (lighter colour) and the model with additional non-obligate animals (darker colour) are shown together. The immigration rate was set to *µ* = 0.1.

### 3.3 Trait space is covered faster in the presence of non-obligate animal mutualists, but the final network structure remains highly specialized

The assembly simulations with only obligate mutualists start from one interacting pair and expand from there into the adjacent trait space. The assembly process in the presence of non-obligate animals, on the other hand, allows plants (and subsequently animals) to immigrate at multiple points in trait space already at early assembly stages, as can be seen in Fig. 3 when comparing the second and third row with the first row.

**Figure 3:**
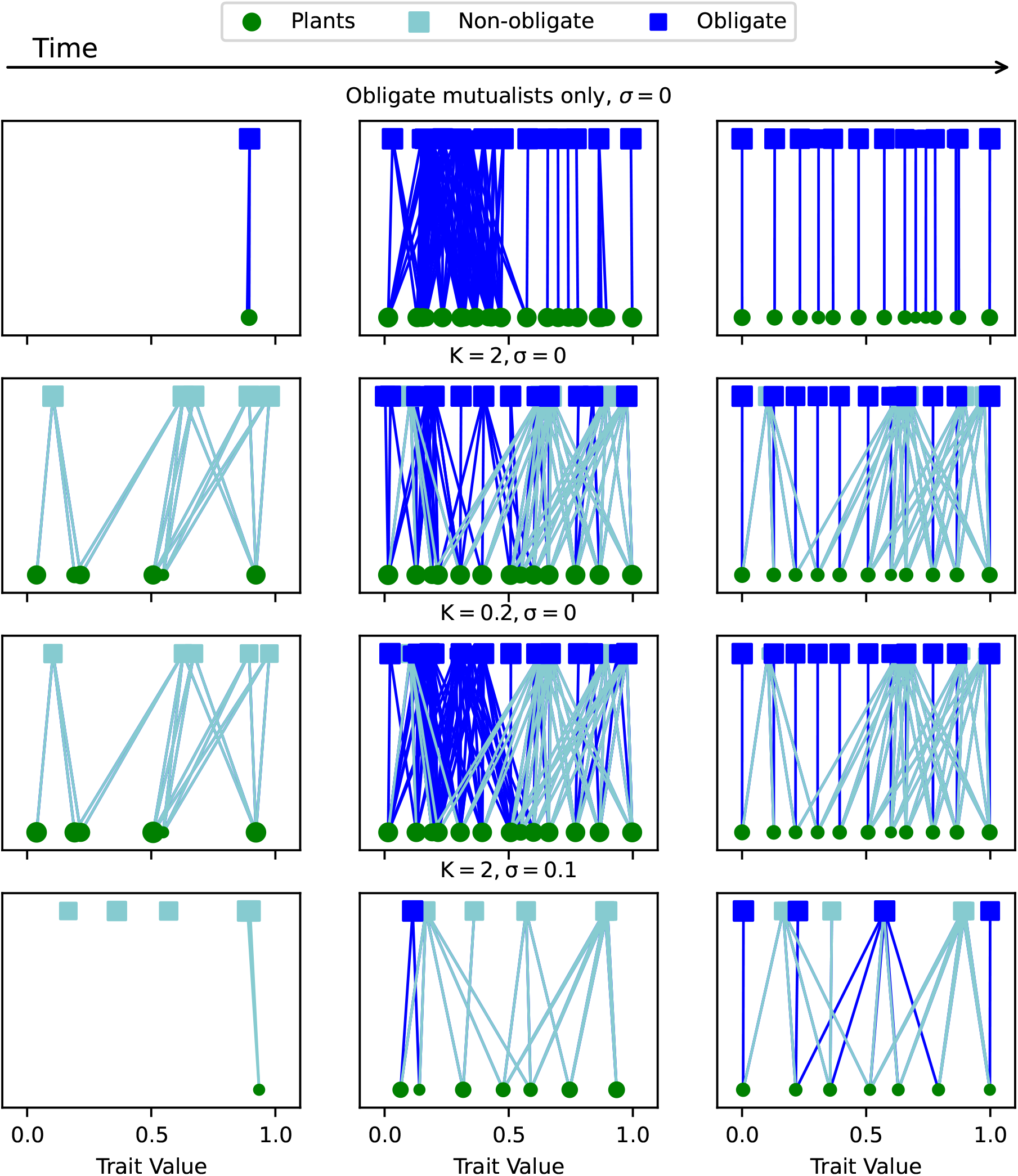
Snapshots of the network assembly after 5 (left), 500 (middle), and 500,000 (right) immigration events. We compare the model with only obligate animals (top row) with the one with non-obligate animals (three bottom rows), the model with deterministic population dynamics (*σ* = 0) to the one with demographic noise (*σ* = 0.1), and simulations with high (*K* = 2) and low (*K* = 0.2) carrying capacity of non-obligate animals. Light blue squares correspond to non-obligate animal mutualists, dark blue to obligate animal mutualists. The circles represent plants. The size of squares and circles corresponds to the population density of the respective species.

Even in the presence of non-obligate animals, the simulation leads to a specialized obligate network, with each plant and each obligate animal having one link. The only animal generalists present at the end are none-obligate animals, as their composition does not change according to our model rules. Again, the population densities of non-obligate animal mutualists has no clearly visible effect on the findings. In the presence of demographic noise, species numbers are smaller, and at the final stage of the assembly obligate generalist and specialist species can coexist. In contrast to the deterministic case, some plants are only connected to a non-obligate animal.

### 3.4 The main reproductive service of animals to plants shifts during assembly from generalist to specialist and from non-obligate to obligate animals

At early assembly stages, plants depend fully on the non-obligate animals for reproduction, and the reproductive service (defined in Eq. (10)) provided by these mutualists is initially 100% (see Fig. 4). Among them, the generalists with large niche width (0.225 *< s*_*j*_ *<* 0.275) contribute considerably more to pollination than the specialists with small niche width (0 *< s*_*j*_ *<* 0.05). With time, the reproductive service of non-obligate animals decreases asymptotically towards a smaller value, which is zero in the simulations with deterministic population dynamics and non-zero with demographic noise. When the carrying capacity of the non-obligate animals is higher, the decrease of their reproductive service is slower, and the asymptotic value reached in the presence of demographic noise is higher. This is not surprising as population sizes of non-obligate animals are higher when their carrying capacity is higher, and so is their contribution to the mutualistic service.

**Figure 4:**
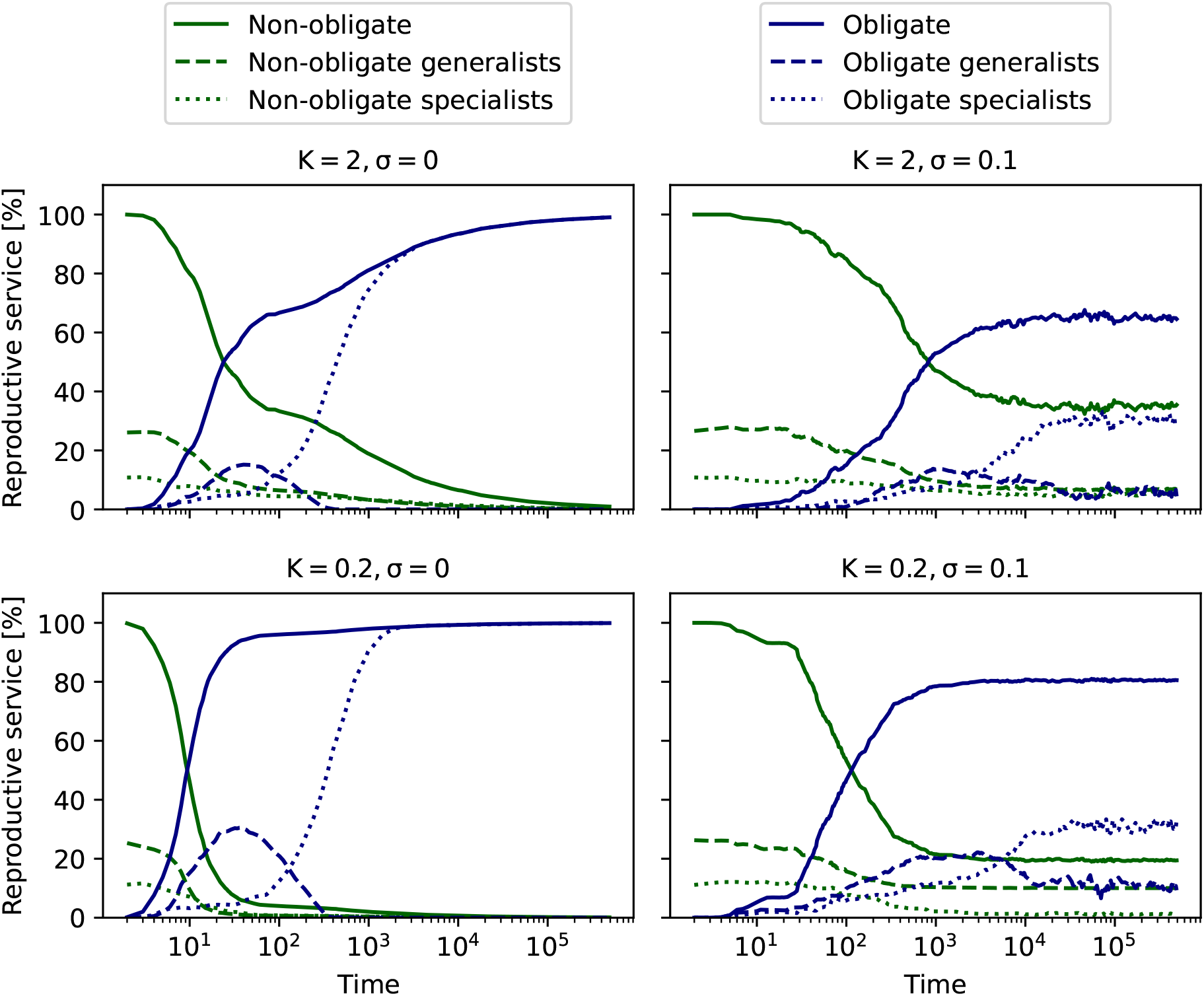
Reproductive service of obligate and non-obligate animal mutualists over time according to Eq. (10). Species with a niche width *s*_*j*_ *∈* [0.225, 0.275] or *s*_*j*_ *∈* (0, 0.05] were labeled generalists (dashed line) and specialists (dotted line), respectively. Full lines correspond to total non-obligate/obligate reproductive service, where the reproductive service of generalist, specialist and intermediate (*s*_*j*_ *∈* [0.05, 0.225]) animal mutualists is added up.

Over time, when the reproductive service by non-obligate animal mutualists decreases, generalist obligate animal mutualists show a peak in reproductive service at intermediate stages, but are gradually replaced by specialist obligate animal mutualists. These specialist obligate animal mutualists provide the major part of reproductive service in the long term in the case with deterministic population dynamics. However, with demographic noise, the reproductive service provided by specialist obligate animal mutualists becomes lower, at a value of approximately 40% for our parameter values. These trends are similar with high and low carrying capacities of non-obligate animal mutualists.

## 4 Discussion

We investigated the assembly of service-for-resource mutualistic networks, using a mathematical model where interactions between species are based on trait matching between plants and animals. Animals in the model may depend exclusively on their plant partners (i.e., they are obligate mutualists), or may have other resources not included in the model (i.e., they are non-obligate mutualists), and both types of animals may be trait specialists that interact strongly with plants within a narrow trait range or trait generalists that interact moderately with plants within a wider trait interval. Our most important findings are that (i) assembly is more rapid if non-obligate animals are present than without them, and (ii) there is a change in species traits during assembly towards a higher degree of specialization, with non-obligate and generalist animals providing the main reproductive service during early stages, and obligate and specialist animals during the late stages. Non-obligatism and generalism thus act in combination to facilitate network assembly. When stochastic effects due to demographic noise are taken into account, the contribution of non-obligate and generalist obligate animals to reproductive service remains non-zero even for long times.

The insights provided by our model have broad implications as the model is not restricted to a specific type of mutualism or animal group, and as the non-obligate animals can be viewed either as facultative mutualists (e.g. flies, butterflies and wasps in pollination mutualism) that rely on resources not included in the model, or as non-resident animals (e.g. mobile pollinating or seed-dispersing birds, bats or insects) that occasionally visit the investigated site but also depend on structures and resources in the surrounding landscape. The observed facilitation effect due to non-obligate animals is supported by empirical observations which report habitat and diet generalist seed-dispersing birds (which correspond to non-obligate animals in our model) to promote tropical forest regeneration Carlo & Morales (2016). Our model suggests that this keystone role of non-obligate animals at early assembly stages is not restricted to plant-seed disperser mutualism, but a feature of any type of service-for-resource mutualism.

The shift in traits of animal mutualists over time observed with our model (see Fig. 4) agrees with general insights of succession theory that early pioneer species differ in their properties from climax species (Chazdon, 2014). Habitat or diet generalists prevail in young forests, while forest specialists or obligate animal mutualists (i.e. diet specialists) dominate in mature forests (Bowman *et al*., 1990; May, 1982; Pardini *et al*., 2009; Pinotti *et al*., 2015). In contrast to our model, each of these studies focuses on specific animal groups (e.g. birds and butterflies), or on a certain type of mutualism (e.g. seed dispersal). Furthermore, the studies consider different types of generalism, namely habitat generalism, diet generalism (when, e.g., a bird eats insects and nectar), or trait generalism (when, e.g., bees visit a broad range of flowers). Our theoretical approach connects these different observations by suggesting that the shift from generalists to specialists occurs in a correlated manner for all three concepts of generalism.

Since the facilitation of assembly by non-obligate animals is similarly enhanced in the case with high and with low population density of non-obligate animals, our model suggests that non-obligate animal mutualist population density is not important for facilitating the network assembly. This finding appears empirically relevant as one can expect the population density of animals in recovering sites to be relatively low in the beginning, due to a lower abundance of resources (Chazdon, 2014).

We could ascribe the facilitation effect to a significantly increased immigration success of plants and animals (Fig. 2). A closer look at snapshots of the network assembly in Fig. 3 revealed that assembly with non-obligate animals can proceed at multiple places in trait space from the beginning, compared to an initially narrow trait range in the absence of non-obligate animals. This means that a higher fraction of immigrating species can establish a link to a species already present in the network, and hence can grow and survive. Additionally, in the network with only obligate species newly entering species are necessarily ecologically similar to species already present in the network, which means that their growth is hampered by competition for resources and for mutualists. By enabling immigration of plants over a wide range of traits, non-obligate animals lead to decreased interspecific competitive pressure but have a competitive disadvantage themselves, and they are replaced by obligate animals during succession.

Immigration success of animals is also enhanced, despite the presence of non-obligate animals with similar traits. We explain this with the relatively low competition between obligate and non-obligate animals in our model. As we assume food to be the limiting resource for animals, non-obligate animals can always rely on their other nutrition source apart from the one provided by their interaction partners in the recovering habitat, leaving more of these resources to the obligate animals. In this way, the competition between obligate animal mutualists is decreased because they share less plant partners in the early stage compared to the model version with only obligate species.

One might argue that the increased immigration success in the presence of non-obligate animals is due to a wider range of trait space being available, increasing the chance of an immigrating species to establish a link. To test this hypothesis, we also made a simulation where each immigrating species is chosen such that it has at least one link upon entry into the network. We found that in this case immigration success is still higher in the model version with non-obligate animal mutualists than without (see Figure S1 in Supporting Information). This means that the availability of niches alone cannot fully explain the increased immigration success, but the reduced competition also plays an important role.

Non-obligate animals lead in our model to functional redundancy and therefore to a more robust network structure during later assembly stages: whenever the obligate partner animal of a plant goes extinct, the non-obligate partner might save it. Higher robustness due to functional redundancy is supported by empirical observations (Biggs *et al*., 2020). With demographic noise, the non-obligate animal mutualists provide a non-negligible proportion of reproductive service even in the long term, due to the continuous extinction risk of obligate animals and the subsequent necessity of plants to rely on their non-obligate partners. Hence non-obligate partners stabilize the mutualism against reproductive failure and co-extinction.

In conclusion, our model provides a unifying framework and a mechanistic understanding of different empirical observations concerning the role of diet, habitat, and trait generalists during early stages of ecosystem recovery and the shift towards a higher degree of specialism at later stages. It demonstrates that these shifts occur in combination, suggesting a broader empirical exploration of pioneer and climax species’ traits to corroborate and refine our mechanistic findings. Ultimately, this can show ways for accelerating ecosystem restoration.

## Acknowledgements

This research is part of the DFG Research Unit FOR 5207 “Reassembly of species interaction networks”. We thank the members of this Research Unit for helpful discussions, and David Donoso, Carsten Dormann, and Jochen Fründ for helpful comments on an earlier version of the manuscript. We thank Nico Blüthgen, David Donoso, Maria-José Endara, Constance Tremlett and Karin Römer for project coordination and administration. We acknowledge financial support from the DFG via grant number DR300-17.

---

**Table 1:** List of all parameters and variables that were used in the simulations, including a description, the used values and respective dimensions.

| Var. | Description | Value | Dimension |
| --- | --- | --- | --- |
| $t_{P_i}$ | Plant niche value | $\in [0,1]$ | |
| $t_{A_j}$ | Animal niche value | $\in [0,1]$ | |
| $s_j$ | Animal niche width | $\in (0,0.275]$ | |
| $\xi$ | Plant niche width | 0.1 | |
| $p_i$ | Density of plant population i | | [individuals/area] |
| $a_j$ | Density of animal population j | | [individuals/area] |
| $r$ | Per capita growth rate of plants | 1 | [1/time] |
| $\lambda$ | Per capita growth rate of animals | 0.8 | [1/time] |
| $w$ | Intraspecific competition coeff. of plants | 0.8 | [area/individuals] |
| $u$ | Interspecific competition coeff. of plants | 0.2 | [area/individuals] |
| $\mu_p$ | Per capita mortality rate of plants | 0.002 | [1/time] |
| $\mu_a$ | Per capita mortality rate of animals | 0.1 | [1/time] |
| $K$ | Carrying capacity | 0.2, 2 | [area/individuals] |
| $\beta$ | Logistic growth rate | 2 | [1/time] |
| $\sigma$ | Strength of demographic noise | 0, 0.1 | $[\sqrt{\frac{\text{individuals}}{(\text{area} \cdot \text{time})}}]$ |
| $\mu$ | Immigration rate | 0.1 | [1/time] |
| $\theta_p$ | Extinction threshold of plants | $10^{-2}$ | [individuals/area] |
| $\theta_a$ | Extinction threshold of animals | $10^{-4}$ | [individuals/area] |
| | Initial population density of plants | $(1+10^{-4}) \cdot \theta_p$ | [individuals/area] |
| | Initial population density of animals | $(1+10^{-4}) \cdot \theta_a$ | [individuals/area] |
| $dt$ | Step size for numerical integration | 0.1 | [time] |

