## Supplementary File 1 for "Shifts from non-obligate generalists to obligate specialists in simulations of mutualistic network assembly"

### Appendix S1

In our model, traits of immigrating plants and animals were chosen randomly between 0 and 1 upon immigration. If only obligate species are present, immigration success is small during early stages of the assembly since only a small part of the trait space is covered by species. The immigration success was considerably larger when the simulations were performed in the presence of non-obligate animals, see Fig. 2. To identify whether the increased chance to establish a link is the sole reason for this enhanced immigration success, we investigated a model variant where we allowed only those plants and animals to invade that have a link to a plant or animal already present. This enhances immigration success for plants and animals in particular in the case with only obligate species because plants and animals without links are not considered at all for immigration. Fig. S1 shows that immigration success of plants and animals is still higher at the early stage of the assembly in the case with non-obligate animals compared to the case with only obligate animals.

However, in the case with only obligate animals the immigration success has now increased compared to the simulations shown in Fig. 2, where no constraint was imposed on the possible trait values of species that immigrate. This means that the availability of niche space does not fully explain the difference between the immigration success with non-obligate animals compared to the case with only obligate animals. We ascribe the remaining difference to reduced competition between immigrating species in the case with non-obligate animals as they can have larger differences in traits during the early stages of the assembly.

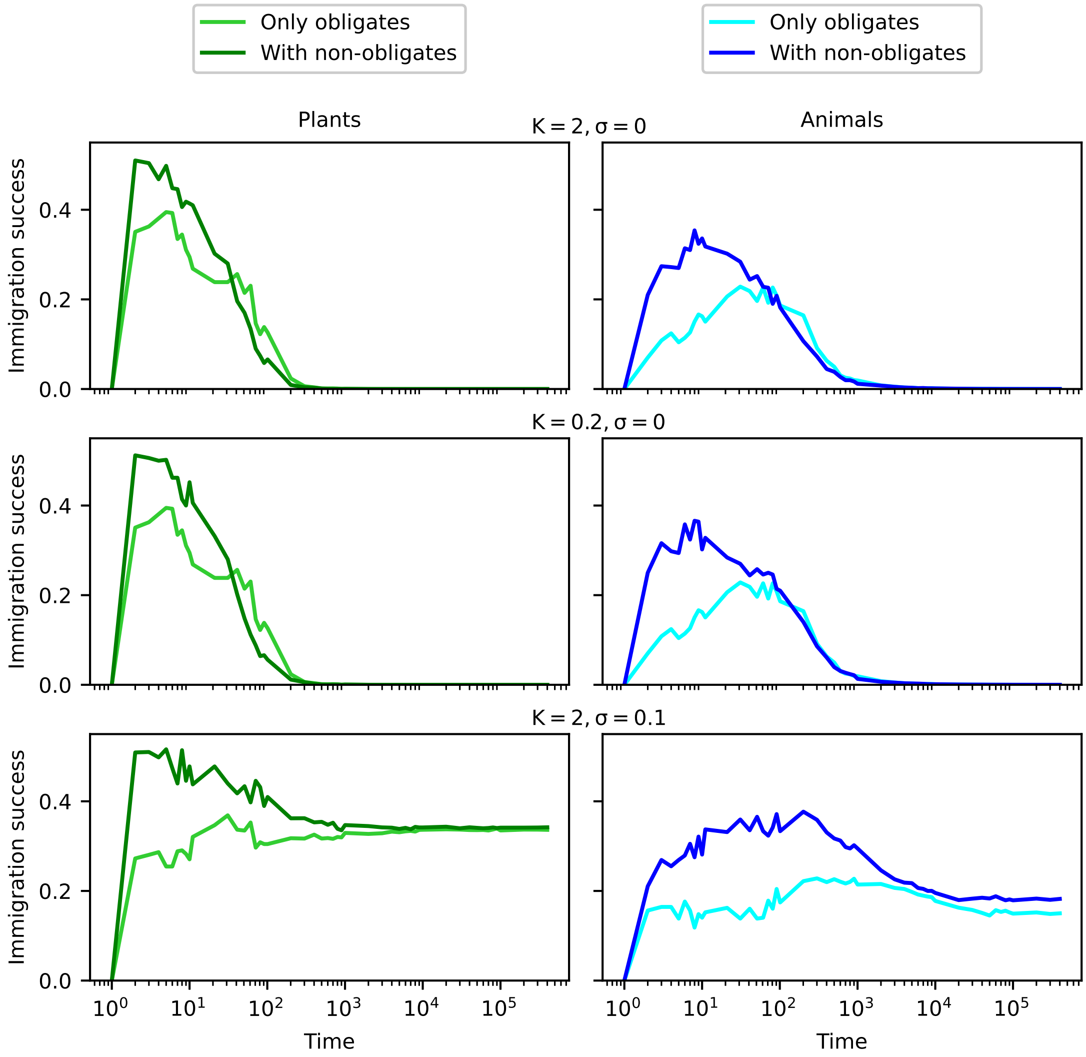


Figure S1: Immigration success of plants (left) and animals (right) over time for the model variant where only those species are chosen for immigration that can establish a link to an existing species. We compare deterministic population dynamics ($\sigma = 0$) and population dynamics with demographic noise ($\sigma = 0.1$), and high ($K = 2$) and low ($K = 0.2$) carrying capacity of non-obligate animals. Computer simulations of the model with only obligate animals (lighter colour) and the model with additional non-obligate animals (darker colour) are shown together. The immigration rate was set to $\mu = 0.1$.
